# Widespread methane formation by *Cyanobacteria* in aquatic and terrestrial ecosystems

**DOI:** 10.1101/398958

**Authors:** M. Bižić-Ionescu, T. Klintzsch, D. Ionescu, M. Y. Hindiyeh, M. Günthel, A.M. Muro-Pastor, W. Eckert, F. Keppler, H-P Grossart

## Abstract

Evidence is accumulating to challenge the paradigm that biogenic methanogenesis, traditionally considered a strictly anerobic process, is exclusive to *Archaea*. Here we demonstrate that Cyanobacteria living in marine, freshwater and terrestrial environments produce methane at substantial rates under light and dark oxic and anoxic conditions, forming a link between light driven primary productivity and methane production in globally relevant group of phototrophs. Biogenic methane production was enhanced during oxygenic photosynthesis and directly attributed to the cyanobacteria by applying stable isotope labelling techniques. We suggest that formation of methane by *Cyanobacteria* may contribute to methane accumulation in oxygen-saturated surface waters of marine and freshwater ecosystems. Moreover, in these environments, cyanobacterial blooms already do, and might further occur more frequently during future global warming and thus have a direct feedback on climate change. We further highlight that cyanobacterial methane production not only affects recent and future global methane budgets, but also has implications for inferences on Earth’s methane budget for the last 3.5 billion years, when this phylum is thought to have first evolved.

## Introduction

Methane (CH_4_) is the second most important anthropogenic greenhouse gas after CO_2_ and is estimated to have 28-34 times higher warming effect than the latter over a 100-year period (Intergovernmental Panel on Climate Change, 2014). The mixing ratio of CH_4_ in the troposphere has increased from 715 ppbv in the preindustrial era to currently 1,860 ppbv (Nov. 2017 NOAA). Estimated global CH_4_ emissions to the atmosphere average at ca. 560 Tg per year (1Tg = 10^12^ g) exceeding the current estimated sinks by ca. 13 Tg per year (Tian et al., 2016). Thus, to mitigate the constant increase in atmospheric CH_4_ a comprehensive understanding of global CH_4_ sources and the environmental parameters that affect them is necessary.

Traditionally, biogenic methanogenesis is the formation of methane under strictly anoxic conditions by microbes from the domain *Archaea* (phylogenetically distinct from both eukaryotes and *Bacteria)*. However, in the past decade there has been growing evidence that also eukaryotes such as algae (Lenhart et al., 2016), plants (Keppler, Hamilton, Braß, & Röckmann, 2006), animals (Tuboly et al., 2013), fungi (Lenhart et al., 2012) and probably humans (Keppler et al., 2016) produce methane under oxic conditions albeit at considerably lower rates. These recent findings suggest that the phenomenon may not be solely limited to methanogenic *Archaea* and could include new metabolic pathways. For example, the conversion of methylated substrates such as methylphosphonates to CH_4_ by *Bacteria* has been extensively addressed in recent years with regards to the “methane paradox” (Repeta et al., 2016; Wang, Dore, & McDermott, 2017). Recently, Zheng *et al*. (2018) have shown CH_4_ formation by *Rhodopseudomonas palustris* during N_2_ fixation. Methane emission was also detected from cryptogamic covers, i.e. phototrophic assemblages on plant, rock and soil surfaces (Lenhart et al., 2015).

Accumulation of CH_4_ in oxygenated freshwater environments has been repeatedly associated with the presence of *Cyanobacteria* (see also Fig. S1). Methane production by *Cyanobacteria* has been attributed to either demethylation of methylphosphonates (Beversdorf, White, Björkman, Letelier, & Karl, 2010; Gomez-Garcia, Davison, Blain-Hartnung, Grossman, & Bhaya, 2011; Yao, Henny, & Maresca, 2016) or to the association of filamentous *Cyanobacteria* with methanogenic *Archaea*, providing the latter with the necessary hydrogen for CH_4_ production (Berg, Lindblad, & Svensson, 2014). *Cyanobacteria* are ubiquitous, found literally in any illuminated environment as well as unexpectedly in some dark subsurface ones as well (Hubalek et al., 2016; Puente-Sánchez et al., 2018). Furthermore, this phylum predominated Earth whilst the environment was still reductive and ca. 1.3 billion years prior to the great oxidation event which occurred 2.4 billion years ago (Gumsley et al., 2017). Therefore, we tested whether this phylum contributes to the global CH_4_ budget independent of naturally occurring, extra-cellular precursor substances or the presence of methanogenic *Archaea*. If so, their ubiquitous nature, their expected future increase in abundance (Huisman et al., 2018; Visser et al., 2016) and their proximity to the interface with the atmosphere makes them potential key players in the global CH_4_ cycle.

Here we demonstrate that unicellular as well as filamentous, freshwater, marine and terrestrial members of the prominent and ubiquitous phylum *Cyanobacteria*, a member of the domain *Bacteria*, produce CH_4_ at substantial rates under both light and dark, oxic and anoxic conditions.

## Material and Methods

### Cyanobacterial cultures

Seventeen different cyanobacterial cultures were obtained from various sources and grown using the media described in Table 1.

**Table 1.**
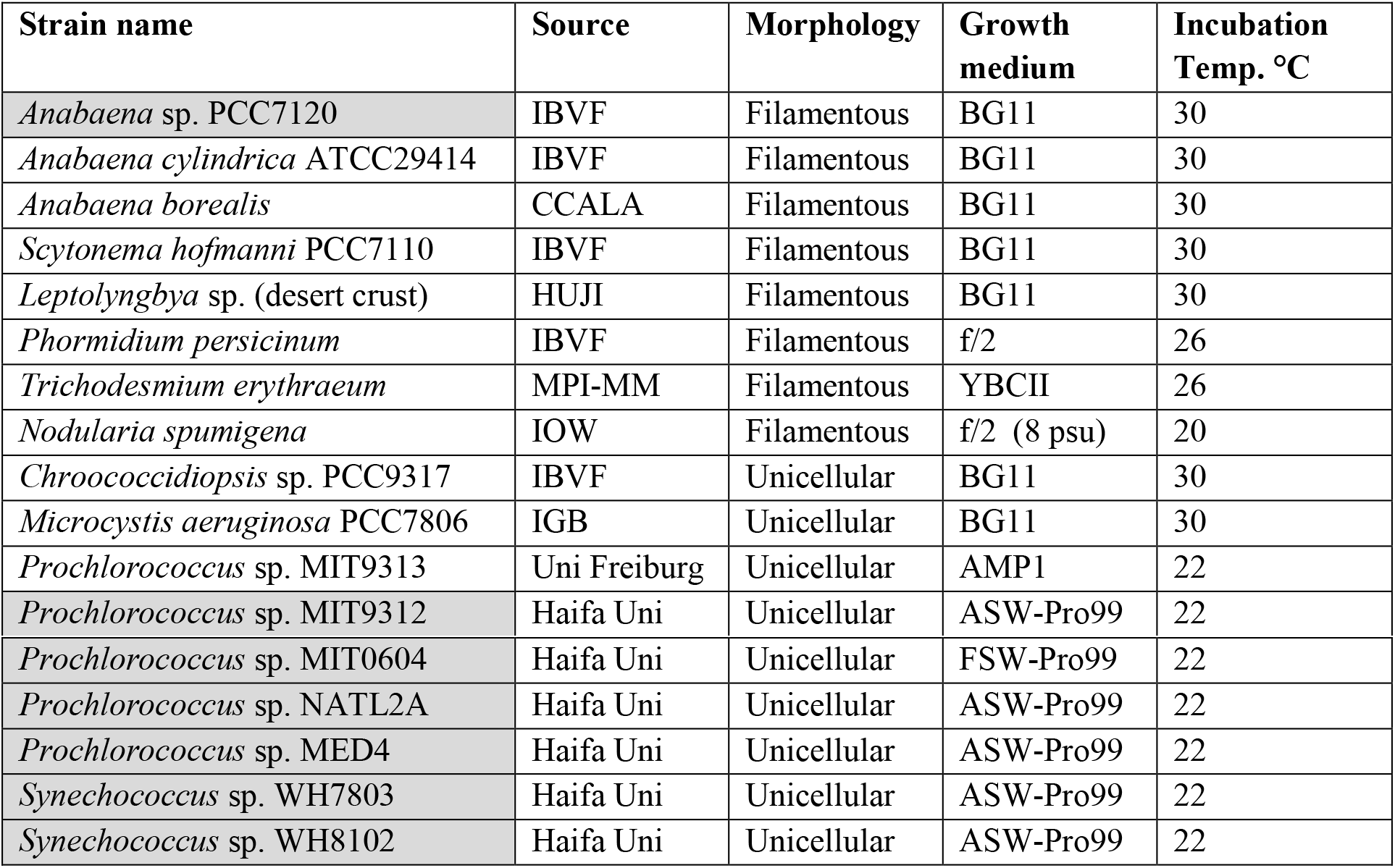
Cyanobacterial cultures used in this study and their growth conditions. Shaded cultures are fully axenic while others are mono-cyanobacterial. Sources abbreviations: IBVF: Culture collection of the Institute for Plant Biochemistry and Photosynthesis, Sevilla Spain; CCALA: Culture collection of autotrophic organisms; HUJI: Laboratory of Aaron Kaplan, Hebrew University of Jerusalem, Jerusalem Israel; IOW: Laboratory of Falk Pollehne, Leibniz Institute for Baltic Sea research, Warnemünde, Germany; MPI-MM: Max Planck Institute for Marine Microbiology, Bremen, Germany; IGB: Leibniz Institute of Freshwater Ecology and Inland Fisheries, Neuglobsow, Germany; Uni. Freiburg, Laboratory of Claudia Steglich, Freiburg University, Freiburg, Germany. Haifa University, Laboratory of Daniel Sher. Media source: BG11 (Stanier, Deruelles, Rippka, Herdman, & Waterbury, 1979); f/2 (Guillard & Ryhter, 1962); YBCII (Chen, Zehr, & Mellon, 1996); AMP1 (Moore et al., 2007); Filtered sea water (FSW) / Artificial sea water Pro99 (Moore et al., 2007).

| Strain name | Source | Morphology | Growth medium | Incubation Temp. °C |
| --- | --- | --- | --- | --- |
| *Anabaena* sp. PCC7120 | IBVF | Filamentous | BG11 | 30 |
| *Anabaena cylindrica* ATCC29414 | IBVF | Filamentous | BG11 | 30 |
| *Anabaena borealis* | CCALA | Filamentous | BG11 | 30 |
| *Scytonema hofmanni* PCC7110 | IBVF | Filamentous | BG11 | 30 |
| *Leptolyngbya* sp. (desert crust) | HUJI | Filamentous | BG11 | 30 |
| *Phormidium persicinum* | IBVF | Filamentous | f/2 | 26 |
| *Trichodesmium erythraeum* | MPI-MM | Filamentous | YBCII | 26 |
| *Nodularia spumigena* | IOW | Filamentous | f/2 (8 psu) | 20 |
| *Chroococcidiopsis* sp. PCC9317 | IBVF | Unicellular | BG11 | 30 |
| *Microcystis aeruginosa* PCC7806 | IGB | Unicellular | BG11 | 30 |
| *Prochlorococcus* sp. MIT9313 | Uni Freiburg | Unicellular | AMP1 | 22 |
| *Prochlorococcus* sp. MIT9312 | Haifa Uni | Unicellular | ASW-Pro99 | 22 |
| *Prochlorococcus* sp. MIT0604 | Haifa Uni | Unicellular | FSW-Pro99 | 22 |
| *Prochlorococcus* sp. NATL2A | Haifa Uni | Unicellular | ASW-Pro99 | 22 |
| *Prochlorococcus* sp. MED4 | Haifa Uni | Unicellular | ASW-Pro99 | 22 |
| *Synechococcus* sp. WH7803 | Haifa Uni | Unicellular | ASW-Pro99 | 22 |
| *Synechococcus* sp. WH8102 | Haifa Uni | Unicellular | ASW-Pro99 | 22 |

### Stable isotope labeling experiments Culturing and treatments

To investigate the production of *Cyanobacteria*-derived CH_4_, 60 ml vials with 40 ml liquid and 20 ml head space volume (ambient air) were used and sealed with septa suitable for gas sampling. For the ^13^C labelling experiments NaH^13^CO_3_ (99 % purity, Sigma-Aldrich, Germany) was added amounting to 10 % of the initial dissolved inorganic carbon (DIC) in BG11 (Stanier et al., 1979) (DIC = 0.4 mM, enriched by added NaHCO_3_; pH ≈ 7.0) and 4.5 % of the DIC in f/2 medium (Guillard & Ryhter, 1962) (DIC = 2015 μmol L^−1^; pH ≈ 8.2) and 1 % of the DIC in the Pro99 (Moore et al., 2007) based medium used for axenic *Synechococcus* and *Prochlorococcus* cultures. Four different examination groups were used: (1) Sterile medium; (2) Sterile medium with NaH^13^CO_3_; (3) Sterile medium with culture; (4) Sterile medium with culture and NaH^13^CO_3_; Four replicates of each cyanobacteria culture (n = 4).The cultures were grown under a light–dark cycle of 16 and 8 hours at 22.5 °C at a light intensity of ≈ 30 μmol quanta m^−2^ s^−1^ for a total period of 3 days.

### Continuous-flow isotope ratio mass spectrometry (CF-IRMS)

CF-IRMS was employed for measurement of the δ^13^C-CH_4_ values in the head space gas above the cultures. Head space gas from exetainers was transferred to an evacuated sample loop (40 mL) and interfering compounds were then separated by GC and CH_4_ trapped on Hayesep D. The sample was then transferred to the IRMS system (ThermoFinniganDeltaplus XL, Thermo Finnigan, Bremen, Germany) via an open split. The working reference gas was carbon dioxide of high purity (carbon dioxide 4.5, Messer Griesheim, Frankfurt, Germany) with a known δ^13^C value of −23.64 ‰ relative to Vienna Pee Dee Belemnite (V-PDB). All δ^13^C-CH_4_ values were corrected using three CH_4_ working standards (isometric instruments, Victoria, Canada) calibrated against IAEA and NIST reference substances. The calibrated δ^13^C-CH_4_ values of the three working standards were −23.9 ± 0.2 ‰, −38.3 ± 0.2 ‰, and −54.5 ± 0.2 ‰. The average standard deviations (n = 3) of the CF-IRMS measurements were in the range of 0.1 to 0.3 ‰. All ^13^C / ^12^C-isotope ratios are expressed in the conventional δ notation in per mille (‰) vs. V-PDB, using the equation 1.

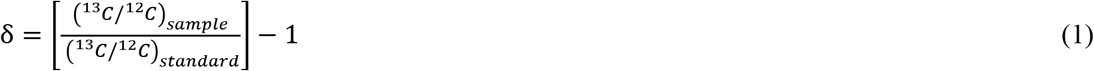

### Membrane inlet mass spectrometer experiments

Experiments were conducted using a Bay Instruments (MD, USA) Membrane Inlet Mass Spectrometer (MIMS) consisting of a Pfeiffer Vacuum HiCube 80 Eco turbo pumping station connected to a QMG 220 M1, PrismaPlus^®^, C-SEM, 1-100 amu, Crossbeam ion source mass spectrometer (Pfeiffer Vacuum, Germany). Culture samples were pumped (Minipuls3, peristaltic pump, Gilson) through a capillary stainless tubing connected to Viton^®^ pump tubing as described in Kana *et al*. (Kana, Cornwell, & Zhong, 2006). The coiled stainless-steel tubing was immersed in a water bath to stabilize the sample temperature. Temperatures were set according to the growth conditions of the different cultures. Inside the vacuum inlet the sample passed through an 8 mm long semipermeable microbore silicone membrane (Silastic^®^, DuPont) before exiting the vacuum and returning to the culture chamber forming a closed system with respect to liquids. This required a 3.5 ml experimental chamber which consisted of an inner chamber where cultures were placed, and an isolated outer chamber connected to a water bath to maintain the culture at a constant temperature. The experimental chamber was placed on a magnetic stirrer and the cultures were continuously stirred for the duration of the experiments to prevent the formation of concentration gradients.

Cultures were transferred to fresh medium before the onset of each experiment after which 3.5 ml of the culture were transferred to the experimental chamber and an equal volume was used for determination of cell counts or dry weight. The latter was determined by filtering the samples on pre-weighed combusted GFF filters (Millipore) and drying at 105 °C for 48 h. In the case of non-homogenous cultures, the biomass from the experimental chamber was used at the end of the experiment for determination of dry weight. Marine picophytoplankton cultures were normalized by cell counting using a FACSAria II flow cytometer (BD Bioscienses, Heidelberg, Germany) at a flow rate of 23.5 μl / min for 2.5 min. Autofluorescence was used to separate cells from salt precipitates in the medium. Cells for counting were collected from the same batch used in the experimental chamber.

The light regime for the experiments was as follows: dark from 19:30 to 09:00 then light intensity was programmed to increase to 60, 120, 180, 400 μmol quanta m^−2^ s^−1^ with a hold time of 1.5 h at each intensity. After maximum light the intensity was programmed to decrease in reverse order with the same hold times until complete darkness again at 19:30. Experiments lasted a minimum of 48 h with at least one replicate longer than 72 h. A minimum of 3 replicate experiments were conducted for each culture.

As negative controls ultrapure water as well as autoclave cyanobacterial biomass were measured to test for non-biogenic methane prosecution by the experimental system (Fig. S2).

### Methane calculations

Methane (and oxygen) concentrations were calculated using the ratio relative to the inert gas Ar (m/z 40). Methane concentration was deduced from mass 15 which does not overlap with other gases in the sample (Schlüter & Gentz, 2008). The CH_4_, O_2_ and Ar concentration in the different media were calculated based on known solubility constants (Powell, 1972) and were calibrated to the measured signals in MQ water and growth media at different temperatures. To further calibrate the CH_4(mz15)_ /Ar ratio, measurements were conducted on air saturated water at different salinities (Fig. S3).

Methane production rates were calculated as the 1^st^ derivative of the Savizky-Golay (Savitzky & Golay, 1964) smoothed data using the sgolay function in the R package signal (http://r-forge.r-project.org/projects/signal/). To account for the continuous degassing from the CH_4_ supersaturation experimental chamber, the degassing rate was determined experimentally using rapid heating of air-saturated water from 18 to 30 °C leading to an instant (super)saturation of 127 % and 130 % for CH_4_ and Ar, respectively. This procedure was repeated under two mixing conditions: I) mixing was applied via magnetic steering as conducted for most cultures; II) mixing occurred only via the cyclic pumping of sample through the MIMS membrane as applied to *Synechococcus* and *Prochlorococcus* cultures. The change in concentration of CH_4_ was monitored and a linear (R^2^ = 0.95) saturation degree dependent rate was determined. The determined rate given in Equations 2 and 3 for type I and type II mixing, respectively, was similar to that determined by comparing the most negative slopes of the culture experiments, when cyanobacterial production rates are expected to be minimal or zero, and the supersaturation state of the culture. Final rates were calculated by adding the absolute values of the local CH_4_ slope (1^st^ derivative) and the local degassing rate (equations. 2,3).

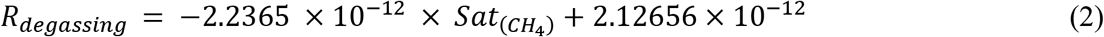

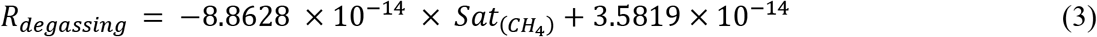

Where: R_degassing_ is the degassing rate in mol CH_4_ sec^−1^ and Sat_(CH4)_ is the fraction CH_4_ saturation state >1 (and <1.3) determined by measured concentration vs. calculated solubility.

### DNA extraction and sequencing

To evaluate the presence of methanogenic *Archaea* in non-axenic cultures DNA was extracted as described in Nercessian *et al*. (Nercessian, Noyes, Kalyuzhnaya, Lidstrom, & Chistoserdova, 2005). The resulting DNA was sent for Illumina sequencing at MrDNA (Shallowater, TX, USA) on a Miseq platform 2×300 bp using the Arch2A519F (CAGCMGCCGCGGTAA) and Arch1017R (GGCCATGCACCWCCTCTC) primers (Fischer, Güllert, Neulinger, Streit, & Schmitz, 2016). Archaeal community composition was analyzed using the SILVA-NGS pipeline (Ionescu et al., 2012) (Fig. S4). After a standard PCR for the *mcrA* gene resulted in no visible products from any of the cultures a qPCR assay was conducted as well resulting in low copy numbers of the gene (Fig. S4). The sequences were submitted to the European Nucleotide Archive under project number: PRJEB25851.

## Results and Discussion

To test the hypothesis that *Cyanobacteria* directly produce CH_4_ independent of methylated precursors (e.g. methylphosphonates) in ambient water, thirteen different filamentous and unicellular cyanobacterial cultures (for details of chosen cultures see Table 1) that are known to grow in marine, freshwater and terrestrial environments were incubated under sterile conditions with ^13^C labelled sodium hydrogen carbonate (NaH^13^CO_3_) as carbon source. All investigated cyanobacterial cultures showed CH_4_ production with increasing stable isotope values (δ^13^C-CH_4_ values) clearly indicating that ^13^C carbon was implemented into CH_4_, whereas no ^13^C enrichment occurred in the control experiments (Fig. 1). These results unambiguously show that *Cyanobacteria* produce CH_4_ *per se* and that the process is most likely linked to general cell metabolism such as photoautotrophic carbon fixation. The different enrichment of ^13^C indicated by δ^13^C-CH_4_ values ranging from 1.71 to 1337 ‰ observed in the different cultures is a result of different production rates as well as differences in biomass. The involvement of methanogenic *Archaea* in this process can be ruled out. First, five of the cultures were axenic. Second, the oxygen concentrations during CH_4_ production were in most cases above saturation level (Fig. 2 and Fig. S2) and while methanogenic *Archaea* were recently reported from oxic environments (Angle et al., 2017), their activity is attributed to anoxic microniches. Third, sequencing analysis of non-axenic cultures and quantitative real-time PCR of the *mcrA* gene showed methanogenic *Archaea* are either absent or present in negligible numbers (Fig. S4).

**Figure 1.**
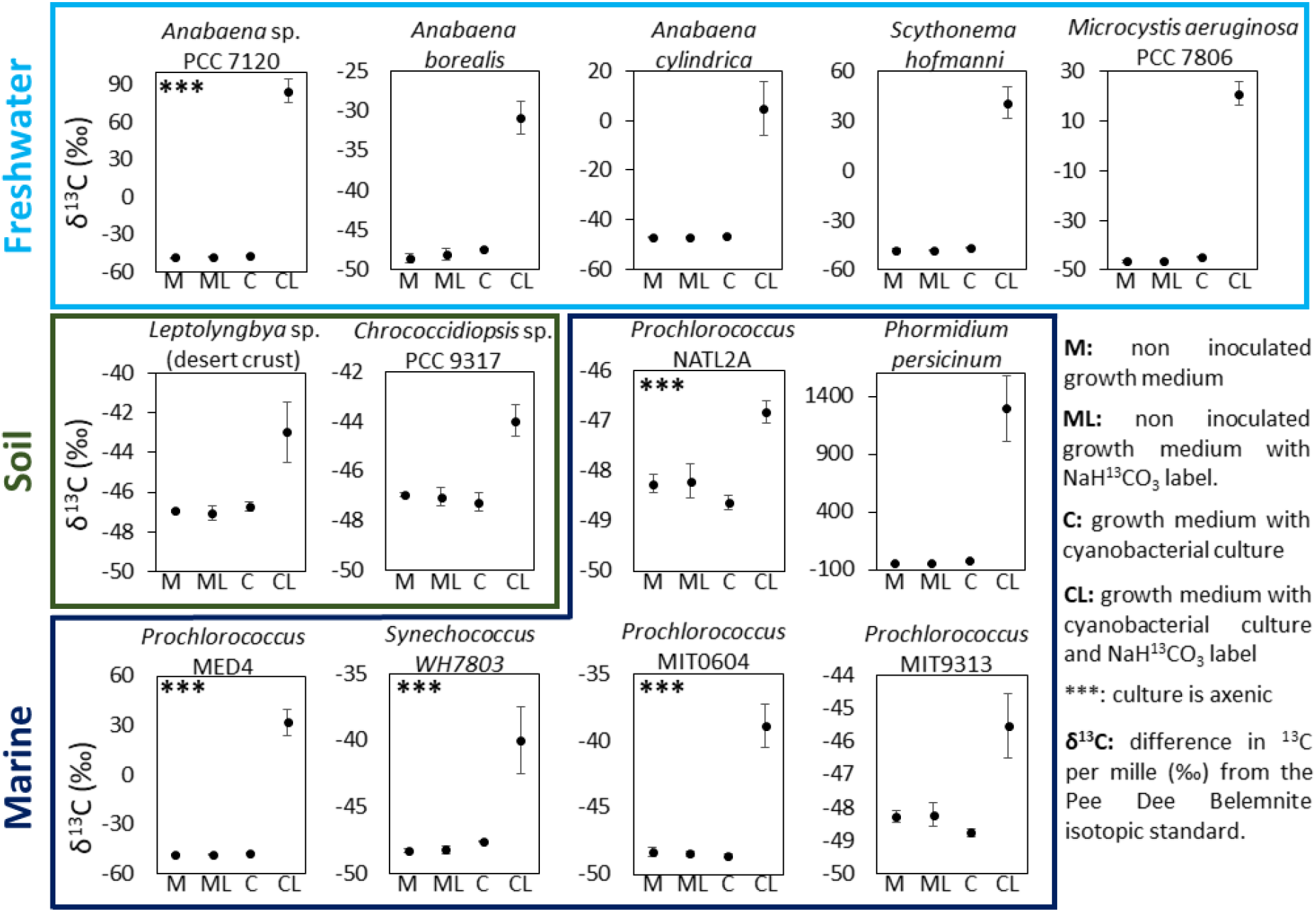
δ^13^C-CH_4_ values measured during incubation experiments of fifteen different filamentous and unicellular freshwater, soil and marine cyanobacterial cultures with and without NaH^13^CO_3_ supplementation. All cyanobacterial cultures produced CH_4_. Using NaH^13^CO_3_ as carbon source (CL) resulted in increasing stable δ^13^C-CH_4_ values as compared to the starting condition. This establishes the direct link between carbon fixation and CH_4_ production. The ^13^C enrichment is not quantitative and thus not comparable between cultures. Error bars represent standard deviation (n=4).

Furthermore, demethylation of methylphosphonates from the spent growth medium is unlikely to be the mechanism involved in this instance even though some *Cyanobacteria* do possess the necessary enzymatic machinery (Beversdorf et al., 2010; Gomez-Garcia et al., 2011) for the following reasons. First, thus far, demethylation of methylphosphonates has been shown to occur only under phosphorus starvation, which was highly unlikely in this study since the culture medium contained ca. 200 μmol P L^−1^. Indeed, publicly available transcriptomic data for *Anabaena* sp. PCC 7120 (Flaherty, Van Nieuwerburgh, Head, & Golden, 2011; Mitschke, Vioque, Haas, Hess, & Muro-Pastor, 2011) and *Trichodesmium erythraeum* (Pfreundt, Kopf, Belkin, Berman-Frank, & Hess, 2014) show no evidence for activity of the phosphonate C-P lyase genes under standard (P-rich) culture conditions. Secondly, some of the *Cyanobacteria* used in this study (i.e. *Microcystis aeruginosa* PCC 7806, *Synechococcus* WH7803 and WH8102, as well as all sequenced species of *Chroococcidiopsis* sp., *Leptolyngbya* sp. and *Phormidium* sp. and *Prochlorococcus*) do not possess the known C-P lyase genes necessary for conversion of methylphosphonates to CH_4_. The lack of the *phn* genes (gene operon for phosphonate metabolism) was demonstrated to be a common feature of the genus *Prochlorococcus* (Luo & Konstantinidis, 2011). *T. erythraeum* was shown to internally produce phosphonates as P storage later to be freed by demethylation (Dyhrman, Benitez-Nelson, Orchard, Haley, & Pellechia, 2009), a process that is likely to release CH_4_. Nevertheless, the same study shows, though not focusing on cyanobacteria alone, that marine unicellular organisms such as *Synechococcus* and *Crocosphaera*, do not contain a detectable phosphonate storage.

Despite the recent finding of CH_4_ production during N_2_ fixation by *Rhodopseudomonas palustris* (Zheng et al., 2018) we suggest that this is not the pathway leading to CH_4_ production in our experiments. First, most *Cyanobacteria* used in this study are unable (i.e marine *Synechococcus, Prochlorococcus, Microcystis aeruginosa*) or unknown (*Leptolyngbya* sp., *Phormidium persicinum*) to fix N_2_. Second, all experiments were conducted in NO_3_^−^ or NH_4_^+^ rich, fresh, media, and therefore N_2_ fixation in capable cyanobacteria is likely to be inhibited to a certain degree (Knapp, 2012). Thus, given the rapid and tight response of CH_4_ production with the onset of light, we consider that the mechanism by which *Cyanobacteria* readily convert fixed CO_2_ to CH_4_ under light conditions must revolve around their central metabolism. Inhibitors of photosynthesis such as Atrazine and DBMIB (2,5-Dibromo-6-isopropyl-3-methyl-1,4-benzoquinone) inhibit the methane production under light conditions, however, the exact biochemical pathway(s) involved in cyanobacteria-derived CH_4_ formation remain so far unknown and thus require further investigation.

Patterns and rates of CH_4_ production were investigated in seventeen cultures over several days of continuous measurement of CH_4_ concentration using a membrane inlet mass spectrometry system (MIMS). Our measurements, lasting 2-5 days, showed that CH_4_ production occurs both under light and dark conditions (Fig. 2 and Fig. S2). This is evident by a positive production rate at almost all times in all experiments. Replicate experiments revealed that, while *Cyanobacteria* repeatedly produced CH_4_, rates and patterns were not consistent, particularly so for production during the periods of darkness. Often, a period with lower rates of CH_4_ production was observed between light and dark phases (Fig. 2 and Fig. S2). The latter is evidenced as a decrease in CH_4_ concentration resulting from degassing of our incubation system. This suggests that different mechanisms may be involved in CH_4_ production under light and dark conditions, presumably dependent on freshly generated photosynthetic products during light and on storage compounds during dark periods. Fermentation of storage compounds by *Cyanobacteria* has been previously described and known to produce among other compounds acetate and hydrogen which are known precursors of acetoclastic CH_4_ formation (Stal & Moezelaar, 1997). Interestingly, most of the genes required for methanogenesis are present in non-methanogenic organisms, including *Cyanobacteria*. Nevertheless, in this instance since the methyl-coenzyme reductase (*mcr*) gene is absent this would suggest that if *Cyanobacteria* produce CH_4_ via conventional pathways, an ortholog of the *mcr* gene exists.

**Figure 2.**
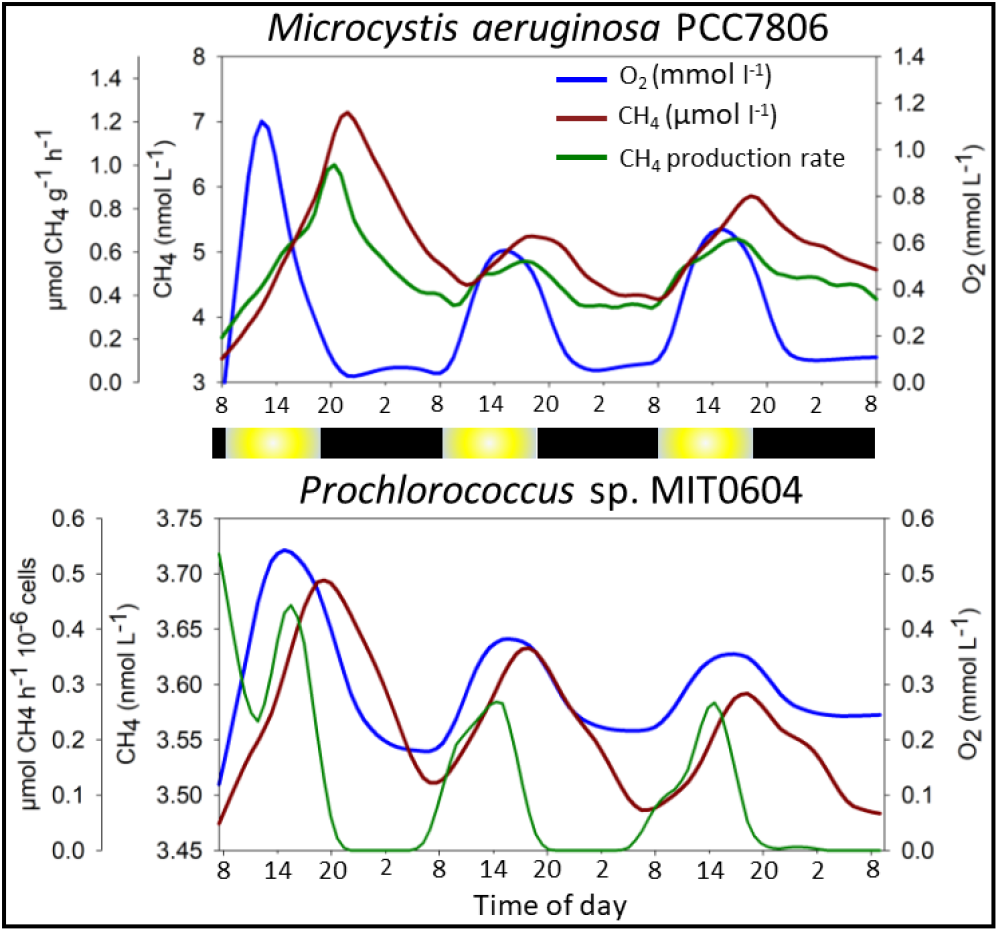
Continuous measurement of CH_4_ and oxygen under light/dark periods using a membrane inlet mass spectrometer (MIMS). Examples are shown for two cultures. Data for other cultures can be found in Fig. S2. A decrease in CH_4_ concentration is a result of either reduced, or no, production coupled with degassing from the supersaturated, continuously-mixing, semi-open incubation chamber towards equilibrium with atmospheric CH_4_ (2.5 nM and 2.1 nM for freshwater and seawater, respectively). Calculated CH_4_ production rates account for the continuous emission of CH_4_ from the incubation chamber for as long as the CH_4_ concentrations are supersaturated. The light regime for the experiments was as follows: dark (black bar) from 19:30 to 09:00 then light intensity (yellow bar) was programmed to increase to 60, 120, 180, 400 μmol quanta m^−2^ s^−1^ with a hold time of 1.5 h at each intensity. After the maximum light period the intensity was programmed to decrease in reverse order with the same hold times until complete darkness again at 19:30.

Methane production rates (Fig. 3) were calculated using the slope of CH_4_ profiles and were normalized on a cyanobacterial biomass dry weight basis for larger cyanobacteria or cell counts for small-celled marine picophytoplankton. The latter to obtain high accuracy for the small-cell-sized picophytoplankton, *Synechococcus* and *Prochlorococcus*. Hourly CH_4_ production rates across cultures of larger cyanobacteria were in the range of 0.1 to 3.4 μmol g^−1^ h^−1^ in individual experiments and a mean of 0.51 ± 0.26 μmol g^−1^ h^−1^. Among the marine picophytoplankton *Synechococcus* sp. exhibited low rates ranging between 10^−4^ and 10^−2^ μmol CH_4_ per 10^6^ cells, while *Prochlorococcus* cultures produced methane at rates ranging from 0.01 to 5 μmol CH_4_ per 10^6^ cells. When compared to production rates of typical methanogenic *Archaea*, CH_4_ production rates of freshwater, soil and large marine cyanobacteria are three to four orders of magnitude lower than the CH_4_ production rates noted for typical methanogenic *Archaea* in culture under optimal conditions (oxygen free) but one to three orders of magnitude higher than rates observed in eukaryotes (Fig. S5). Due to their small size, conversion of CH_4_ production rates of picophytoplankton to μmol g^−1^ h^−1^ results in values exceeding those of methanogenic *Archaea*. Nevertheless, to obtain 1 g of *Prochlorococcus* cells one would need to integrate 0.1-10 m^2^ over a depth of 200 m (Lange et al., 2018) as compared to ca. 20 Kg of soil for methanogenic *Archaea* (assuming 10^9^ cells per g sediment of which 50 % are methanogens). In our experiments, *Prochlorococcus* and *Synechococcus* cultures produced CH_4_ only at light intensities above 20 μmol quanta m^−2^ s^−1^ and therefore, it is likely that only surface *Procholorococcus* and *Syenchococcus* communities contribute to the oceanic CH_4_ flux to the atmosphere.

**Figure 3.**
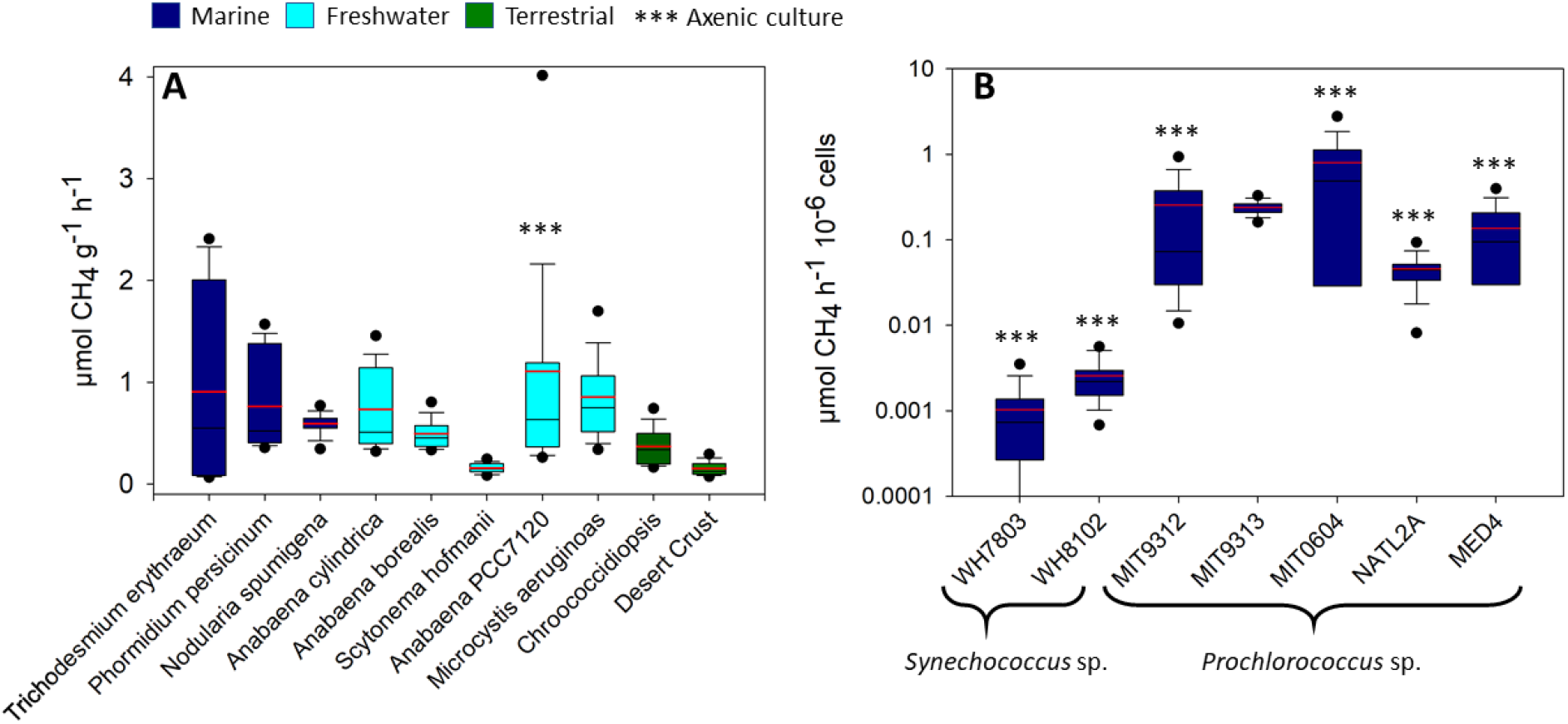
Average CH_4_ production rates (μmol g DW^−1^ h^−1^) obtained from multiple long-term measurements (2-5 days) using a membrane inlet mass spectrometer. The rates are designated by colour according to the environment from which the *Cyanobacteria* were originally isolated; dark blue, light blue and green for marine, freshwater and soil environments, respectively. Black and red lines represent median and mean values, respectively.

Methane production in oxic soils has been previously discussed and attributed mainly to abiotic factors (Jugold et al., 2012) or methanogenic *Archaea* (Hao, Scharffe, Crutzen, & Sanhueza, 1988), although the latter was thought unlikely (Jugold et al., 2012; Kammann, Hepp, Lenhart, & Müller, 2009). Here we show that a typical desert crust *Cyanobacteria* (identified in this study as *Leptolyngbya* sp.), as well as the most common endolithic cyanobacterium *Chroococcidiopsis* (Garcia-Pichel, Belnap, Neuer, & Schanz, 2003) produce CH_4_ both under light and dark conditions (Fig. 1, Fig. S2), thus inferring a new but as yet unknown and unaccounted for source of CH_4_ from oxic soils.

*Cyanobacteria* are ubiquitous in nature and their presence in aquatic systems is expected to increase with eutrophication and rising global temperatures (Visser et al., 2016). The “methane paradox” describing the production of CH_4_ in oxic water layers has been known for four decades (Scranton & Farrington, 1977). Though values may vary between water bodies, a recent study suggests that up to 90 % of CH_4_ emitted from freshwater lakes can be produced in the oxic layer (Donis et al., 2017) with *Cyanobacteria* often being associated with elevated CH_4_ concentration in oxygen supersaturated freshwater systems (Grossart, Frindte, Dziallas, Eckert, & Tang, 2011). In open oceanic environments, distant from any coastal, the contribution of lateral transport from anoxic environments is expected to nonexistent. Nevertheless, based on the emission rates of our laboratory investigations it is difficult to extrapolate the contribution of cyanobacteria to marine, freshwater and terrestrial environments and finally to the global scale. First, only one attempt has been done to estimate the global cyanobacterial biomass (Garcia-Pichel et al., 2003). This study does not account for the increase in blooms of toxic and non-toxic cyanobacteria in freshwater systems (Bowling, Blais, & Sinotte, 2015; Glibert, Maranger, Sobota, & Bouwman, 2014; Huisman et al., 2018; Paerl & Huisman, 2008; Visser et al., 2016), nor for less monitored cyanobacterial environments such under the ice-cover of frozen lakes (Bižić-Ionescu, Amann, & Grossart, 2014). Recent evaluations of *Prochlorococcus* (Lange et al., 2018) suggest a global biomass larger by 33 % than estimated in 2003 by Garcia-Pichel *et al*. Second, while our experiments demonstrate inarguably the ability of *Cyanobacteria* to produce CH_4_ independent of external substrates, as well as to transfer fixed CO_2_ to CH_4_ under laboratory conditions, we cannot account for the effect of nutrients concentrations and light quality in the natural environment. Nevertheless, to get a first idea of what the laboratory rates might sum up to when applied to the natural environment we performed a simple mathematical exercise for the oceanic *Prochlorococcus* community. As suggested before, since experiments with low light (< 20 μmol quanta m^−2^ s^−1^) showed no detectable CH_4_ production only surface *Prochlorococcus* communities were used for this calculation. As such, only rates from High-Light *Prochlorococcus* strains were used i.e. MIT9312, MIT0604 and MED4 averaging at 0.4 μmol CH_4_ h^−1^ 10^−6^ cells. Based on recent estimates of *Prochlorococcus* abundances (Lange et al., 2018) the standing stock global surface communities were estimated to consist of 2.48×10^19^ cells (out of a total of 3.4×10^27^). When taken together, these numbers result in a potential production by global surface *Prochlorococcus* communities of 1.39 Tg CH_4_ y^−1^. This number is not to be confused with the oceanic CH_4_ flux, estimated at 1.2 Tg CH_4_ y^−1^ (Rhee, Kettle, & Andreae, 2009), which is the result of production by multiple processes, transport and consumption by methanotrophs.

In this study, we show that *Cyanobacteria* can readily convert fixed inorganic carbon directly to CH_4_ and emit the potent greenhouse gas under both light and dark conditions. This is in addition to the already established ability of *Cyanobacteria* to produce CH_4_ by the demethylation of methylphosphonates (Beversdorf et al., 2010; Gomez-Garcia et al., 2011). *Cyanobacteria* as a phylum are the most ubiquitous group of organisms on Earth, thriving in most, naturally and artificially, illuminated environments almost regardless of temperatures, salinity and nutrient concentrations. Accordingly, their ability to produce CH_4_ via different pathways, likely related to their surroundings, makes them important to the present and future global CH_4_ cycle and budget. Even more so, as blooms of cyanobacteria are increasing with eutrophication and rising global temperatures (Huisman et al., 2018; Visser et al., 2016). Furthermore, as phototrophic prokaryotes such as *Cyanobacteria* have been inhabiting Earth for more than 3.5 billion years (Falcón, Magallón, & Castillo, 2010; Frei et al., 2016) they may have had a major contribution to Earth’s CH_4_ cycle such as during the great oxygenation event or even earlier when the conditions on Earth were more reductive favoring CH_4_ production.

Further research, however, is needed to elucidate the biochemical pathways of CH_4_ formation in *Cyanobacteria* and fully assess its possible relevance for ecology and the global CH_4_ budget throughout Earth history and how it might change in the future.

## Acknowledgements

*In-situ* probe data was obtained from the LakeLab (www.lakelab.de) as part of the routine monitoring of Lake Stechlin run by the Leibniz Institute of Freshwater Ecology and Inland Fisheries. We thank the technical assistants of department III for making this data available. We thank the members of the MIBI group and particularly Jason Woodhouse for assistance in sampling, analysis and discussion of the data. We thank Tim Urich from Greifswald University for the qPCR analysis. We thank Claudia Steglich and Wolfgang Hess from Freiburg University for the *Prochlorococcus* culture. We thank Falk Pollehne from the Institute of Baltic Sea Research for the *Nodularia* culture. We thank Meri Eichner from the Max Planck Institute for Marine Microbiology for the *Trichodesmium* culture. We thank Daniel Sher and Dalit Roth from Haifa University for the cultures of *Synechococcus* WH7803 and WH8102 and *Prochlorococcus* MIT9312, NATL2A, MIT0604, MED4. Funding was provided to M.B.I., D.I. and H.P.G. via the DFG-Aquameth project (GR1540-21-1), the BMBF-BIBS project (01LC1501G) and the Human Frontiers Science project (HFSP 2039371). F.K. and T.K. were supported by the German Research Foundation (DFG; KE 884/8-2, KE 884/11-1 and KE 884/16-2).

## Author Contributions

M.B.I., T.K., D.I., F.K., H.P.G., conceived the study and designed the experiments; M.B.I., D.I., M.Y.H., M.G., A.M.M.P., W.E., performed MIMS experiments and *in-situ* measurements and analyzed the data; T.K. performed stable isotope measurements and together with F.K. analyzed the data. M.B.I., D.I., performed microbial community data analysis; A.M.M.P. analyzed transcriptomics data. M.B.I., T.K., D.I., M.Y.H., M.G., A.M.M.P., W.E., F.K., H.P.G. discussed the results and wrote the paper.

---

**Fig. S1.**
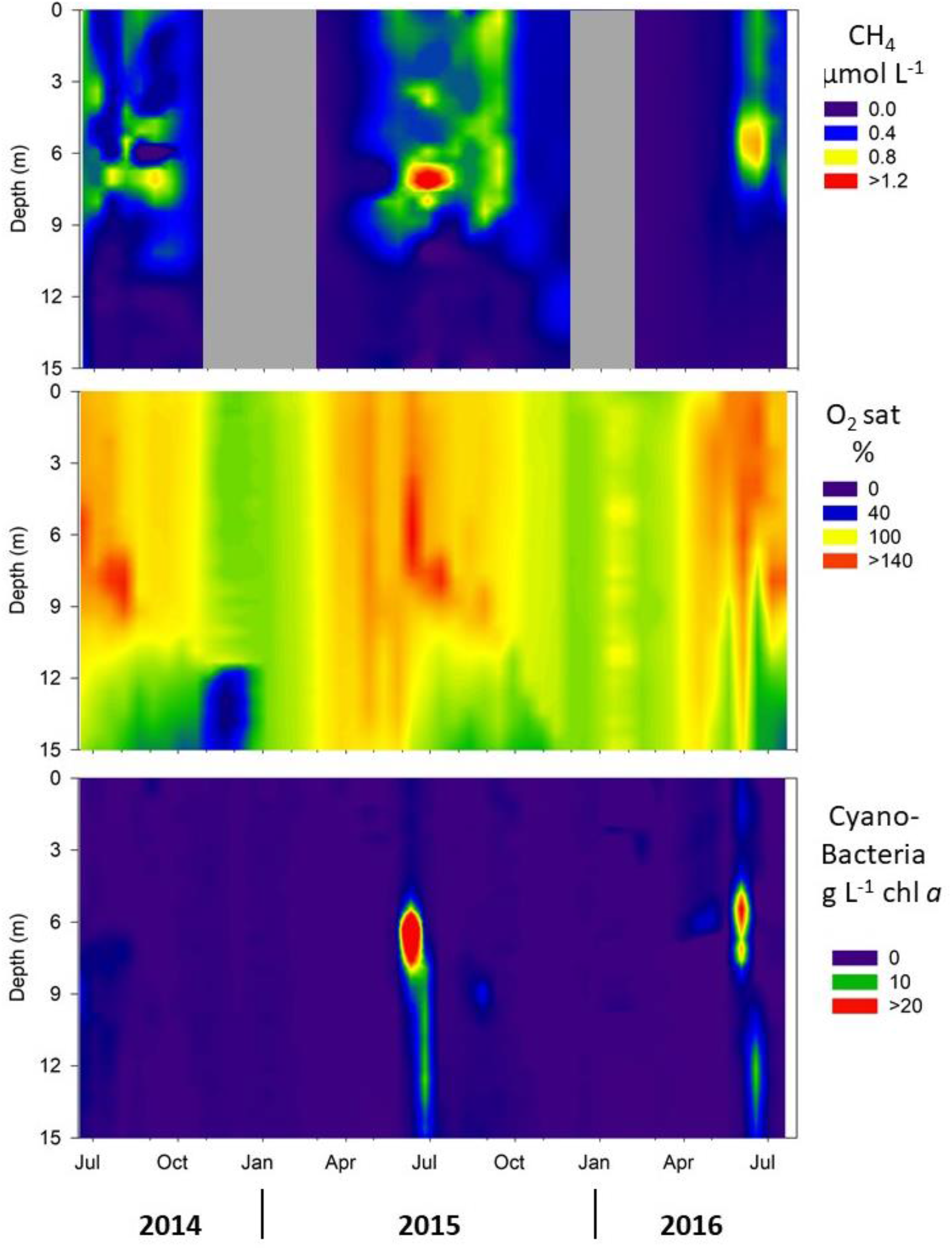
Temporal profiles of CH_4_, O_2_ and cyanobacterial derived Chl a between July 2014 and July 2016. The CH_4_ data was measured every 1 – 4 weeks depending on season using a GC-FID as described in Grossart *et al*. (Grossart *et al*., 2011). O_2_ and Chl a were measured hourly using a YSI and a BBE probe (see www.lake-lab.de), respectively.

**Fig. S2.**
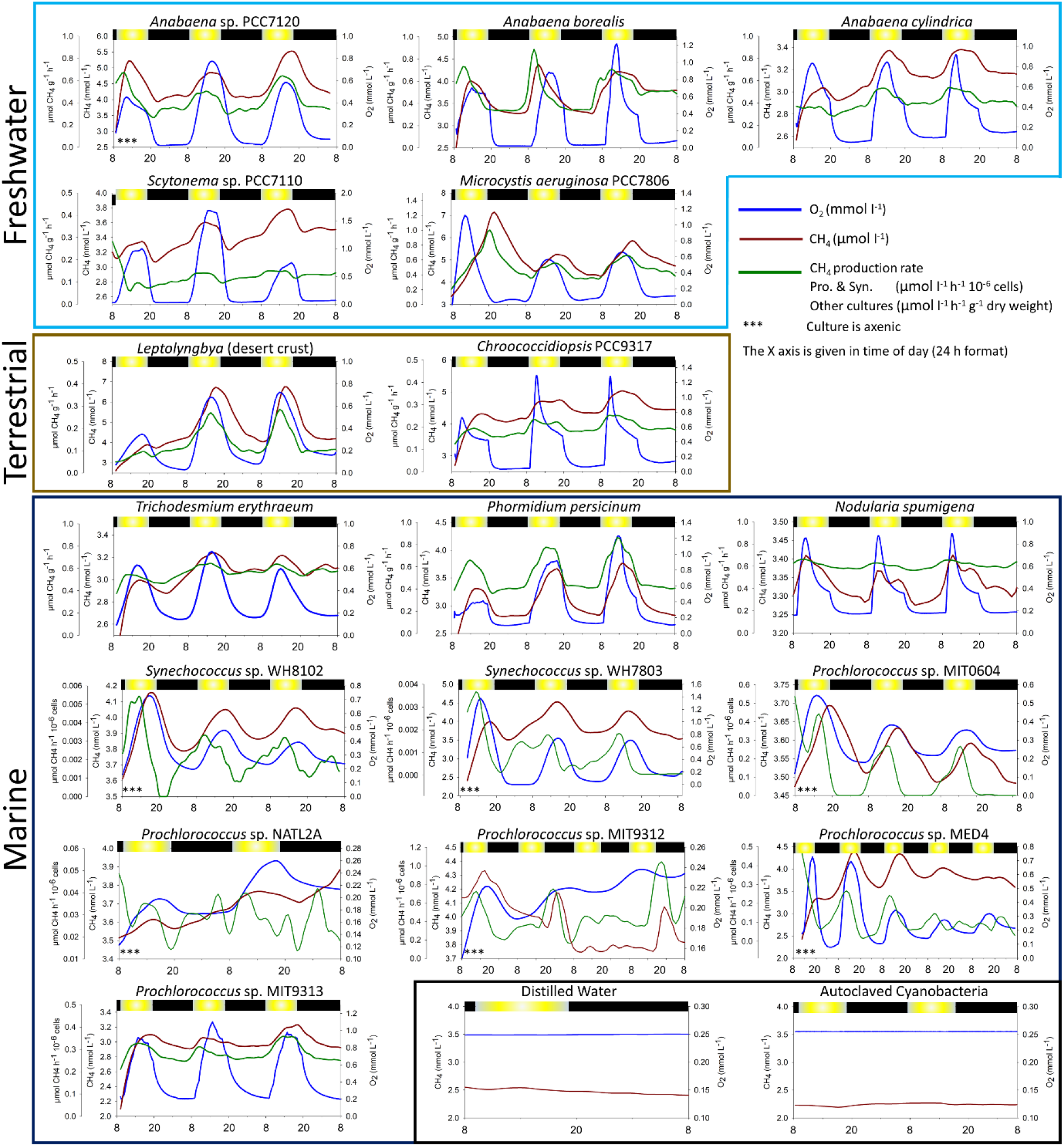
Continuous measurements of CH_4_ and oxygen under light/dark cycles using a membrane inlet mass spectrometer (MIMS) in 17 different cyanobacterial cultures. A decrease in CH_4_ concentration is a result of reduced (or no) production coupled with degassing from the supersaturated, continuously-mixing, semi-open incubation chamber towards equilibrium with atmospheric CH_4_ (2.5 nM and 2.1 nM for freshwater and seawater, respectively). Calculated CH_4_ production rates account for the continuous emission of CH_4_ from the incubation chamber for as long as the CH_4_ concentrations are supersaturated. The light regime for the experiments was as follows: dark (black bar) from 19:30 to 09:00 then light intensity (yellow bar) was programmed to increase to 60, 120, 180, 400 μmol quanta m^−2^ s^−1^ with a hold time of 1.5 h at each intensity. After maximum light period the intensity was programmed to decrease in reverse order with the same hold times until complete darkness again at 19:30.

**Fig. S3.**
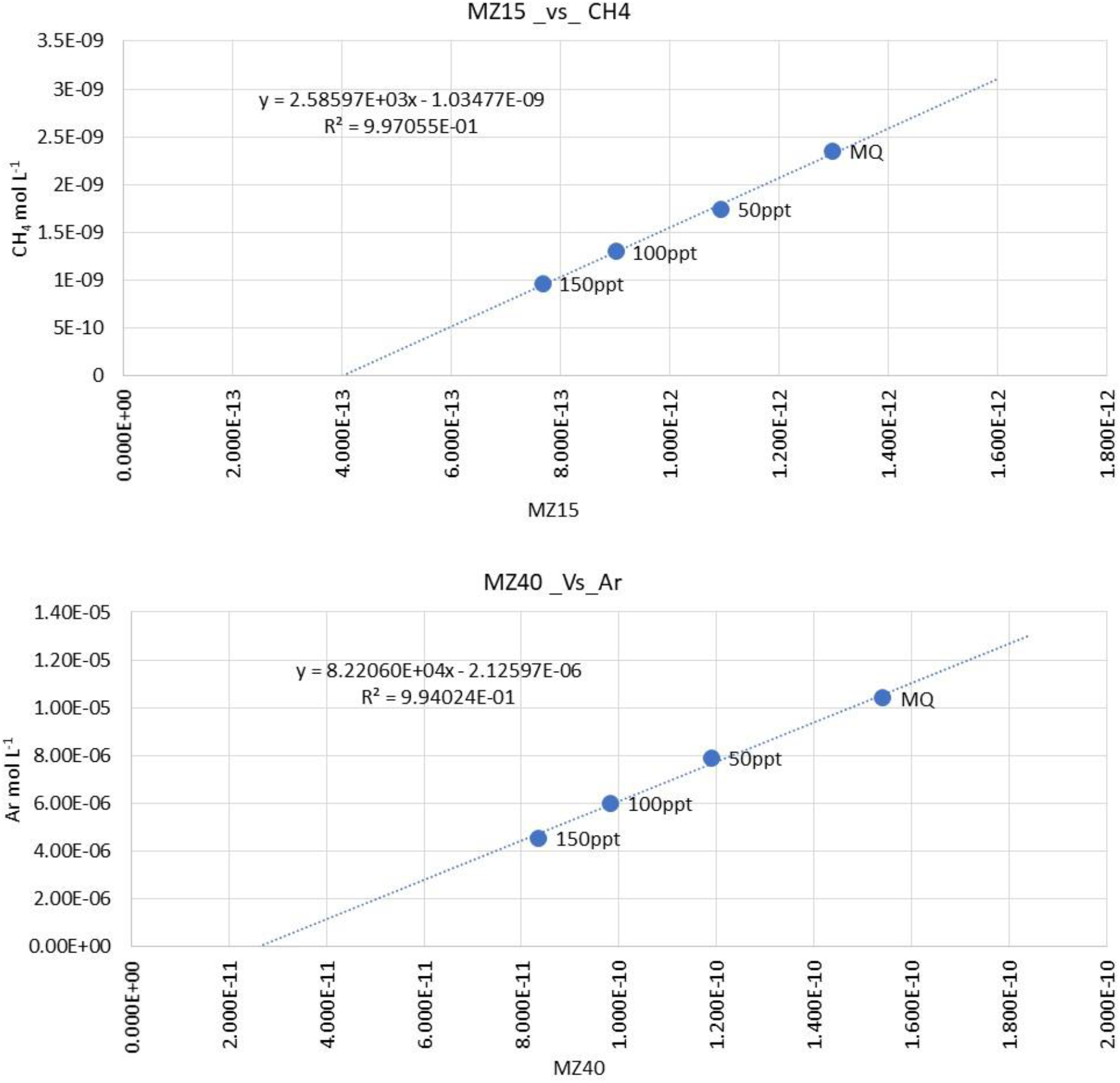
Raw signal obtained from mass 15 (CH_4_) and mass 40 (Argon) plotted against the calculated solubility at different salinities at 30 °C. The signal in both cases is linearly correlated to the concentration of the dissolved gas. The ratio between the two masses was extrapolated between 0 and 50 ppt and was used for calculating the CH_4_ concentration.

**Fig. S4.**
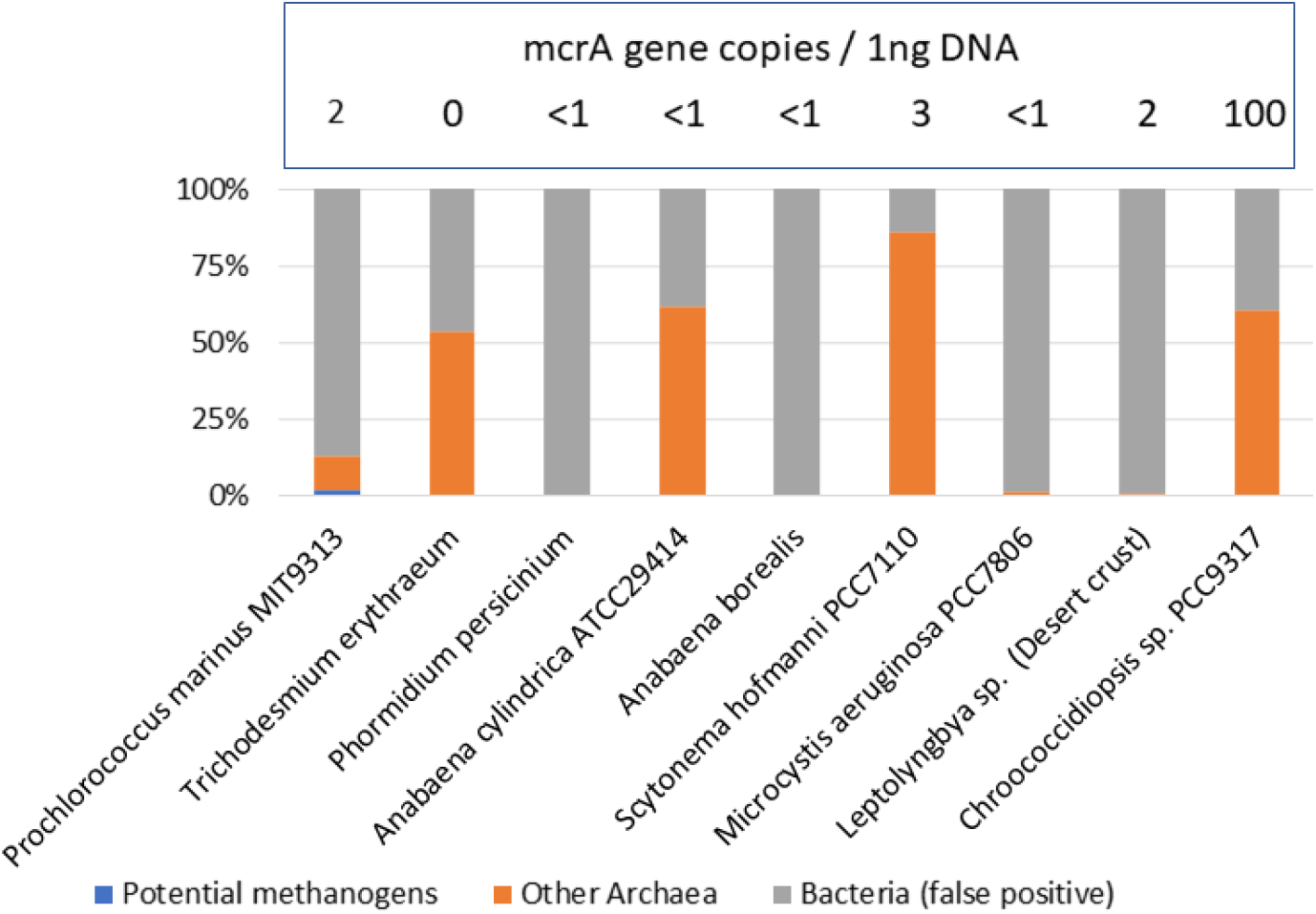
Community composition of the cyanobacterial cultures as obtained when sequenced using *Archaea* specific primers. Methanogenic *Archaea* (red) are in most cases lower that 0.1% of the obtained sequences. In the absence of *Archaea* DNA template, the primers amplify DNA of *Bacteria*. The background presence or complete absence of methanogens was confirmed by qPCR of the *mcrA* gene.

**Fig. S5.**
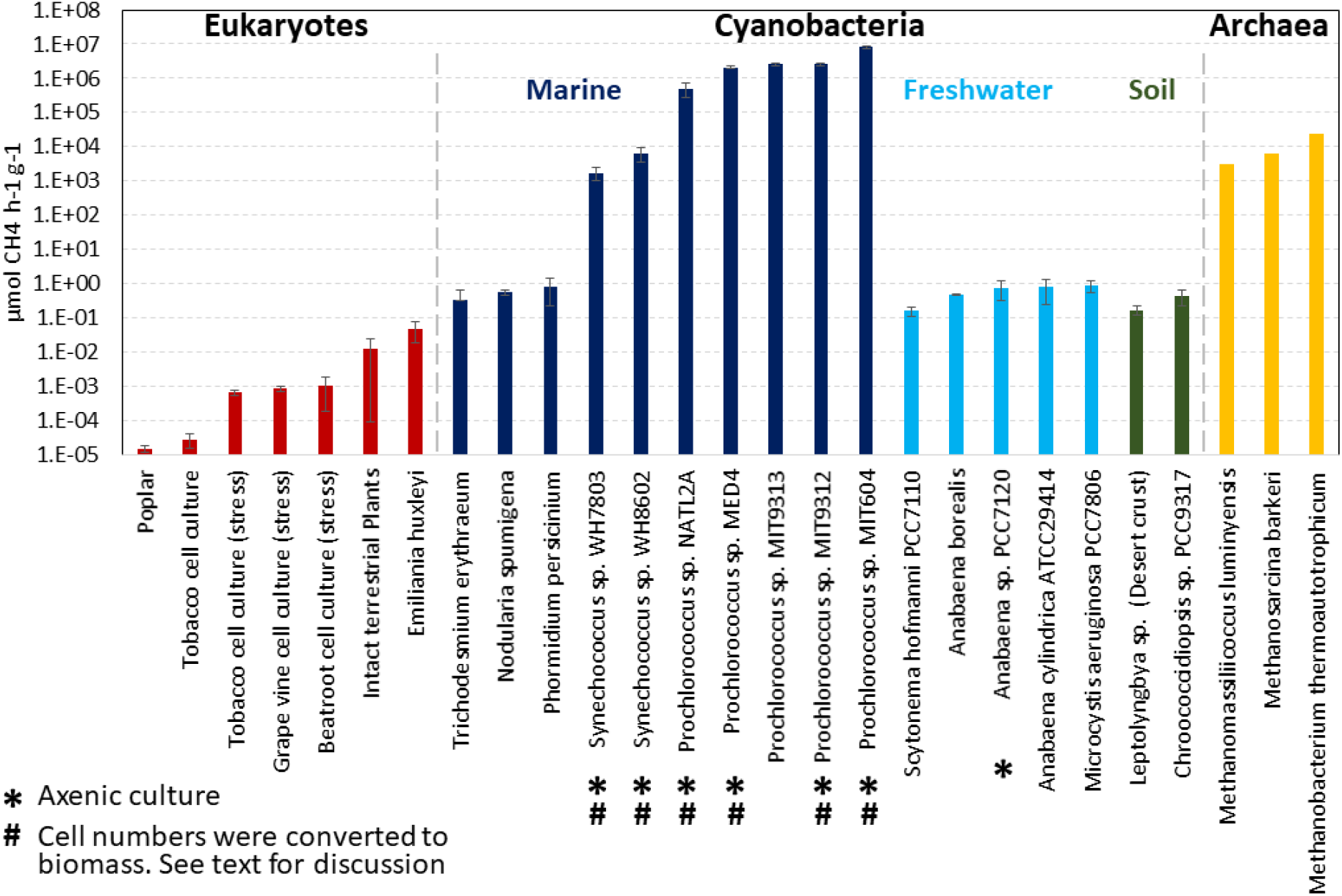
Average CH_4_ production rates (μmol gDW^−1^ h^−1^) obtained from multiple long-term measurements (2-5 days) with a membrane inlet mass spectrometer. The rates are shown by colour according to the environment from which the *Cyanobacteria* were originally isolated; dark blue, light blue and green for marine, freshwater and soil environments, respectively. The rates are presented in comparison to three known methanogens. Rates for the methanogens were obtained from references: Mountfort and Asher (Mountfort and Asher, 1979), Kröninger *et al*. (Kröninger *et al*., 2017) and Gerhard *et* al.(Gerhard *et al*., 1993). Rates for eukaryotes including marine algae and terrestrial plants were taken from Lenhart *et al*. (Lenhart *et al*., 2016), Keppler *et al*. (Keppler *et al*., 2006), Brüggemann *et al*. (Brüggemann *et al*., 2009), Wishkermann *et al*. (Wishkerman *et al*., 2011) and Qaderi *et al*. (Qaderi and Reid, 2009) No emission rates (on a dry weight basis) are available for fungi and animals.

